# Interleukin 22 mediates interleukin 23-induced pathology in newborn mice by disrupting the function of pancreatic and intestinal cells

**DOI:** 10.1101/346577

**Authors:** Glaucia C. Furtado, Lili Chen, Valentina Strohmeier, Zhengxiang He, Madhura Deshpande, Scott K. Durum, Thomas M. Moran, Thomas Kraus, Huabao Xiong, Jeremiah J. Faith, Sergio A. Lira

**Affiliations:** Precision Immunology Institute, Icahn School of Medicine at Mount Sinai, New York, 10029, USA; Institute for Genomics and Multiscale Biology, Icahn School of Medicine at Mount Sinai, New York, 10029, USA; Department of Microbiology, Icahn School of Medicine at Mount Sinai, New York, New York 10029, USA; Center for Therapeutic Antibody Development, Icahn School of Medicine at Mount Sinai, New York, New York 10029, USA; University of Freiburg, Faculty of Biology, Schaenzlestrasse 1, D-79104 Freiburg, Germany; Center for Cancer Research, National Cancer Institute, Frederick, 21702, USA

## Abstract

Mice expressing IL-23 constitutively in the intestine or skin fail to grow and die prematurely. These phenotypes are associated with marked changes in the levels of circulating cytokines and with changes in the transcriptome of the pancreas and intestine. Marked changes are observed in the expression of molecules involved in digestion and absorption of carbohydrates, proteins, and lipids, resulting in a malabsorptive condition. Genetic ablation of IL-22, or one of the subunits of the IL-22R in mice expressing IL-23, restores normal growth and increases the life span of the animals. Mechanistically, IL-22 acts directly at the level of pancreatic acinar cells to decrease expression of the pancreas associated transcription factor 1a (*Ptf1a*), an important transcription factor controlling expression of genes encoding pancreatic enzymes, and acinar cell identity. The results indicate that dysregulated expression of IL-23 and IL-22 has severe consequences in newborns and reveal an unsuspected role for IL-22 in controlling pancreatic enzyme secretion and food absorption.

## Introduction

The gut of a newborn mouse is immature, resembling that of a premature infant(Levy, 2007; Tanner et al., 2015). It lacks cells specialized in antimicrobial defense, such as Paneth cells(Stappenbeck, 2009), contains few T and B cells, and it is populated mostly by myeloid immune cells(Levy, 2007; Marodi, 2006). The lymphoid compartment at this stage is represented mostly by innate lymphoid cells (ILCs) that can respond to bacteria, and are important for formation of structures such as Peyer’s patches and mesenteric lymph nodes(Bar-Ephraim and Mebius, 2016). Maturation of the intestinal barrier and the immune system appear to be dependent of colonization by commensal bacteria(Gensollen et al., 2016; Torow et al., 2017). Colonization of the neonatal intestine starts at birth, with the newly acquired bacteria coming from the vaginal canal or skin, and triggering a complex interaction with cells of the immune system(Gensollen et al., 2016). This process is further influenced by immunoregulatory factors present in the maternal milk(Maynard et al., 2012).

Precise and balanced responses are performed by the cells of the innate immune system in the intestine of newborns, but in a number of occasions, the responses are not balanced, and significant immunopathology arises. One of the important regulators of myeloid and innate lymphoid cell biology is the cytokine interleukin 23 (IL-23)(Basha et al., 2014; Willems et al., 2009). Deficit in IL-23 signaling is not associated with a major deficit in the immune function in newborn mice, but overexpression of IL-23 in neonates has severe consequences, leading to premature death(Chen et al., 2015). Previous work from our group showed that early expression of IL-23 in the neonatal intestine promoted an increase in the number of type 3 innate lymphoid cells (ILC3), altered barrier function and promoted severe intestinal inflammation(Chen et al., 2015). To further understand the pathological consequences of IL-23 expression in early life, we generated two novel mouse strains. We show that expression of IL-23 by CX3CR1-positive cells leads to mortality in 50% of the mice within the first 48 h and that this process is driven by the microbiota, as there is a substantial extension of survival in germ-free conditions. Expression of IL-23 by the keratin-14 promoter also results in premature death, with mice succumbing before 15 days of age. The neonatal mortality associated with IL-23 expression is mediated by its downstream cytokine interleukin 22 (IL-22). IL-22 expression in newborn mice affects expression of genes encoding antimicrobial peptides and digestive enzymes produced by the pancreas and intestine, resulting in a malabsorptive condition. The malabsorptive condition is due in part to downmodulation by IL-22 of the expression of the pancreas associated transcription factor 1a (*Ptf1a*), an important transcription factor required for the maintenance of acinar cell identity and function(Hoang et al., 2016; Walczak et al., 1975). Deletion of IL-22 in mice expressing IL-23 constitutively reverses these changes and prevents early death.

## Results

### Constitutive expression of IL-23 in CX3CR1* cells results in early lethality

Animals expressing IL-23 from the villin promoter (*V23* mice) die at birth(Chen et al., 2015). The cause of death appears related to intestinal bleeding originating from the small intestine. To further study the biology of IL-23 in neonates we engineered mice in which expression of IL-23 was targeted constitutively to CX3CR1^+^ cells, which are the cells that predominantly express IL-23 in the intestine. This was accomplished by intercrossing mice containing a IL-23 cassette preceded by a floxed STOP signal in the ROSA 26 locus (*R23* mice)(Chen et al., 2018), with mice containing a cre recombinase gene inserted into the *CX3CR1* locus (*CX3CR1-cre* mice) (Yona et al., 2013). We refer to these animals as *CXR23* mice (**Fig. 1a**). Expression of the IL-23 subunits p19 and p40 was detected in the intestine of the transgenic, but not control mice (**Fig 1b**). *CXR23* mice were normal at birth, but approximately 50% of the pups died within the first 48 hours of life, with the remaining pups perishing before day 8 (**Fig. 1c**). *CXR23* mice had a normal body weight at birth, but did not grow after birth (**Fig. 1d** and **e**). Necropsy of the newborn mice showed the presence of blood within the small intestine, but not in other organs. Histological analyses of the *CXR23* intestine showed that the bleeding (**Fig 1g**) originated from disrupted villi and from cellular aggregates that resembled PP anlagen (**Fig. 1j**, dashed line) present in the WT small intestine (**Fig. 1i**). The cellular aggregates in the *CXR23* mice were rich in neutrophils that disrupted the overlaying epithelium (**Fig 1h**), and in IL-22^+^ cells (**Fig. 1J**), suggesting a role for these cells in pathology. Of note, the number of ILC3, capable of producing IL-22 and IL17 upon stimulation with IL-23, was markedly increased in the intestine of *CXR23* mice compared to controls (**Fig. 1k**). No abnormalities were found in the large intestine or other organs (kidney, heart, lung and brain) by conventional histological analyses (**Supplemental Figure 1a**). These findings confirm our previous observations that IL-23 expression in the murine gut results in early lethality(Chen et al., 2015).

**Figure 1 -.**
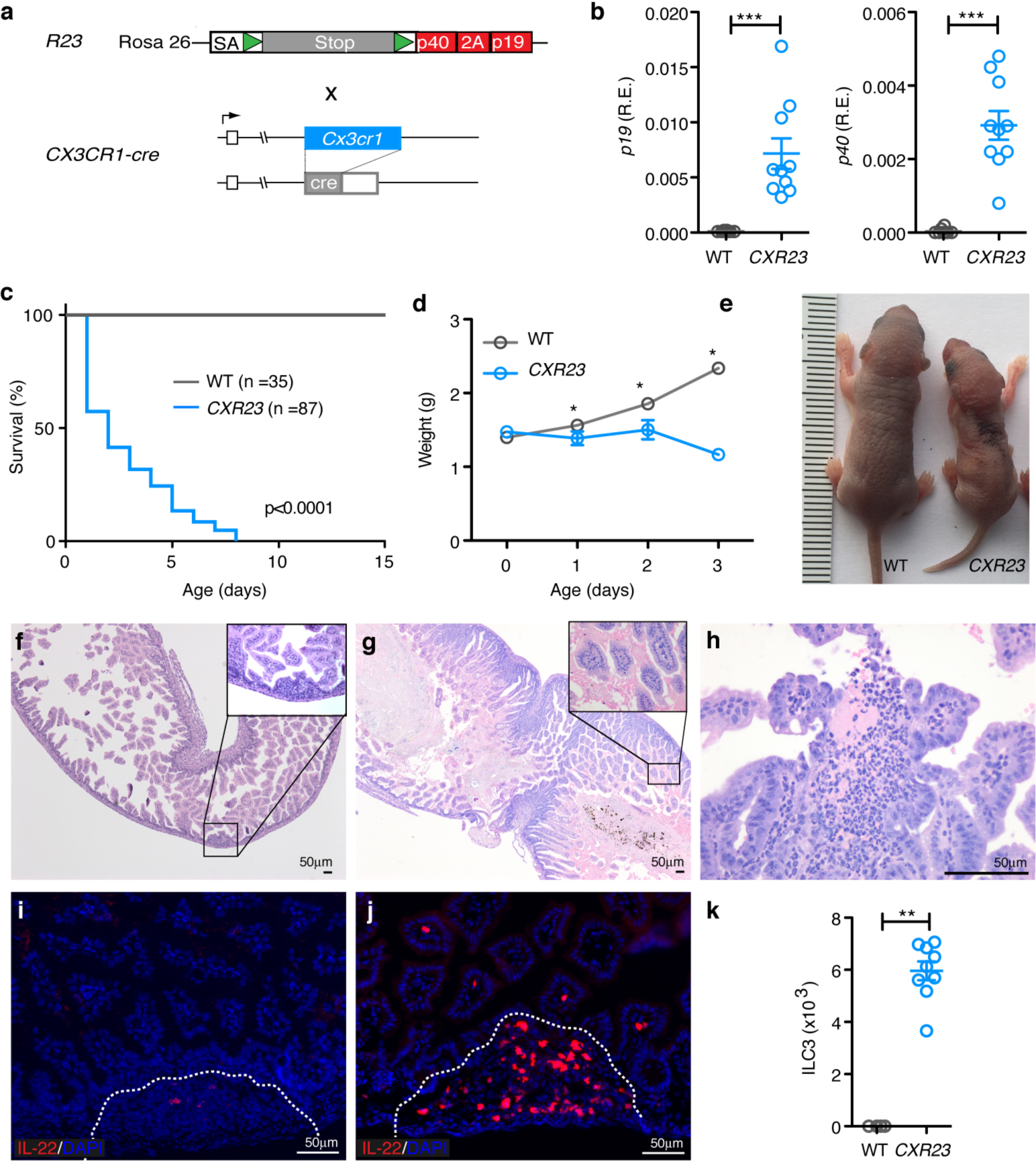
Figure 1 - Constitutive expression of IL-23 in CX3CR1+ cells results in early lethality. (**A**) *R23* mice containing a knockin of IL-23p19 and p40 in the ROSA 26 locus (described in(Chen et al., 2018)) were crossed to *CX3CR1-cre* mice (described in(Yona et al., 2013)) to generate *CXR23* mice. (**B**) Relative expression of *p19* and *p40* mRNA in the intestine of WT and *CXR23* mice at postnatal day 1 (P1) (n = 10 mice/group). (**C**) Survival curves of WT and *CXR23* mice. (**D**) Body weight of WT and *CXR23* mice (n= 11-24 mice/group). (**E**) Representative picture of WT and *CXR23* mice at P3. (F-H) Representative H&E stained section of the small intestine of WT (**F**) and *CXR23* (**G**) mice at P1. Inset shows the presence of red blood cells in the intestine of *CXR23* mice. (**H**) Representative picture of an erosive lesion in the small intestine of *CXR23* mice at P1. (**I**) Immunostaining of the small intestine of *IL-22tdTomato* (**I**) and *CXR23/IL-22tdTomato* (**J**) mice at P1 with anti-tdTomato antibody. Notice the accumulation of IL-22 positive cells in the intestine of *CXR23* mice with erosive lesions. (**K**) Number of ILC3+ cells in the small intestine of *CXR23* mice at P1. Cells were gated on CD45^+^Lin^-^Thy1^+^Sca-1^hi^. Data are shown as mean ± sem, n = 8 mice/group. *p<0.05, **p<0.01, ***p<0.001; by nonparametric Mann-Whitney test.

### *CXR23* germ-free mice have increased lifespan when compared to *CXR23* SPF mice but die at an early age

Previous work from our lab suggested that expression of IL-23 could modify intestinal permeability and facilitate bacterial translocation during the immediate neonatal period(Chen et al., 2015). To investigate whether bacteria contributed to the phenotype of early lethality, we generated *CXR23* mice in germ-free (GF) conditions. Most (>60%) of the *CXR23 GF* mice survived beyond day 10 in contrast to the *CXR23* SPF mice, but succumbed by 30d of age (**Supplemental Fig. 2a**). No bleeding was observed in the intestine of *CXR23 GF* mice at birth or later (**Supplemental Fig. 2b**). Neutrophils were present in the small intestine, but did not disrupt the epithelium (**Supplemental Fig. 2c**). Together, the results suggest that the newly acquired microbiota contributes to the development of the intestinal bleeding phenotype observed in *CXR23* SPF neonates.

### Early lethality in mice expressing IL-23 constitutively in the skin

To further investigate the factors contributing to early lethality and stunted body growth elicited by IL-23 expression, we engineered another transgenic strain in which expression of IL-23 was directed to the keratinocytes, by intercrossing the *R23* mice with mice carrying a cre-recombinase driven by the K14 promoter (**Fig. 2a**). These animals, which we refer to as *KS23* mice, expressed higher levels of IL-23 in the skin than control littermates (**Fig. 2b** and **2c**) and had normal body weight at birth. By day 5 *KSR23* mice had milk in the stomach, but were smaller than their control littermates (**Fig. 2d** and **2e**). Similar to *CXR23* mice, *KSR23* mice died prematurely (before 15 days of age, **Fig. 2f**), but displayed no signs of disease in the skin, intestine, or in other organs examined (**Supplemental Fig. 1b**). A notable finding in the animals was the presence of diarrhea, with yellow stools (**Fig. 2g**), which suggested food malabsorption. Conditions that lead to malabsorption are associated with increased fecal fat content, commonly known as steatorrhea. To determine if *KSR23* mice developed steatorrhea, we measured triglycerides (TG) in the stools and in the serum (**Fig. 2h-i**). At day 5, *KSR23* mice had increased fecal TG (**Fig. 2h**) and decreased TG levels in the serum (**Fig 2i**) when compared to wild type littermate controls. Together, these results indicate that constitutive expression of IL-23 in keratinocytes results in fat malabsorption, growth retardation, and early death.

**Figure 2 -.**
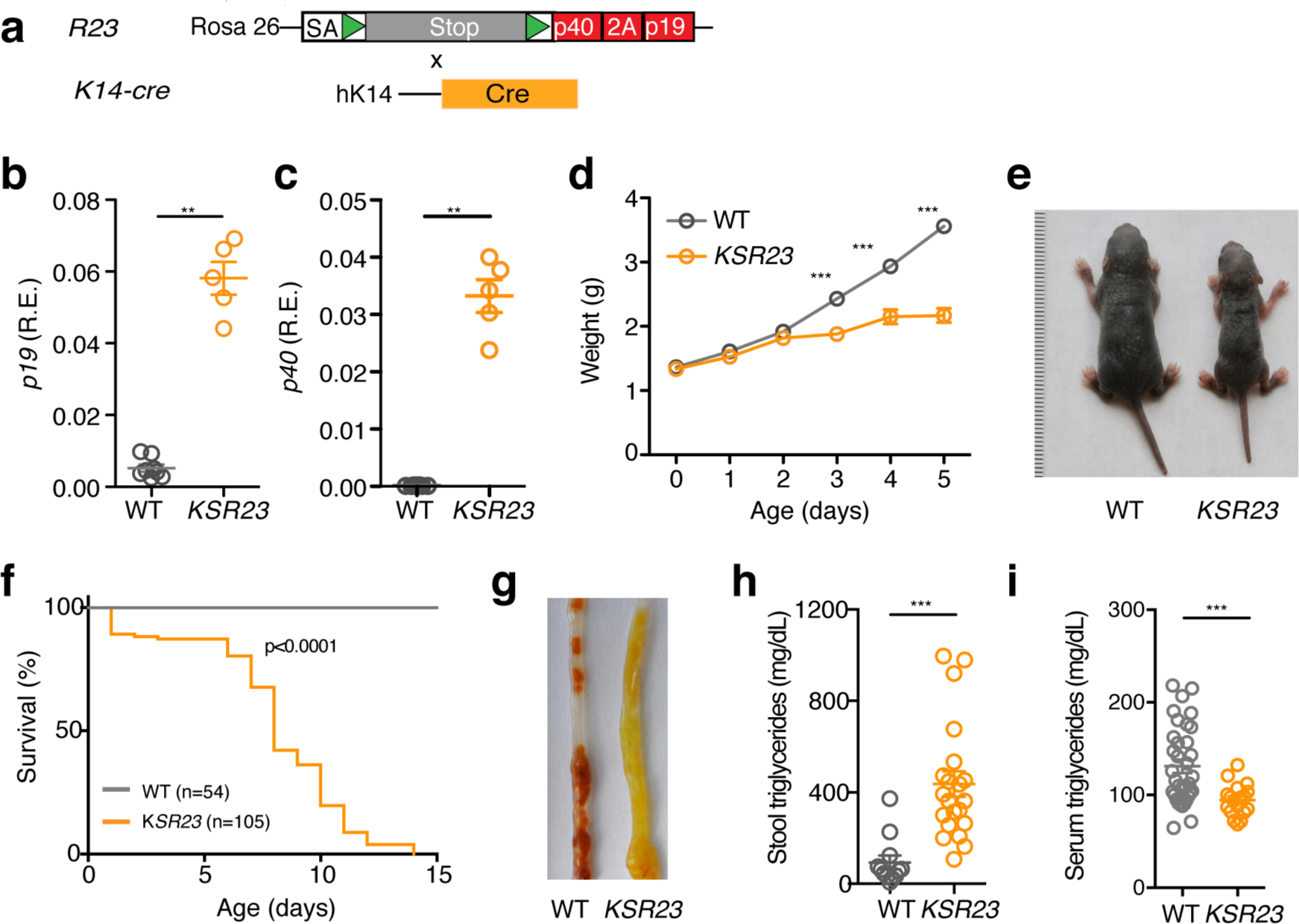
Constitutive expression of IL-23 in K14^+^ cells results in early lethality. (**a**) Scheme for generation of *KSR23* mice. *KSR23* mice were generated by crossing *R23 mice* with mice expressing cre recombinase from the K14 promoter (*K14-cre* mice). Relative expression of (**b**) *p19* and (**c**) *p40* mRNA in the skin of WT and *KSR23* mice at P5 (n = 5 mice/group). (**d**) Survival curves of WT and *KSR23* mice. (**e**) Representative picture of WT and *KSR23* mice at P5. (**f**) Body weight of WT and *KSR23* mice during the first week of life (n= 12-26 mice/group). (**g**) Picture of the large intestine of WT and *KSR23* mice at day 5. Notice the presence of soft yellow stool in the colon of *KSR23* mice. (**h**) Quantification of triglycerides in the stool of WT (n=12) and *KSR23* (n=21) mice at P5. (**i**) Quantification of triglycerides in the serum of WT (n=35) and *KSR23* (n=18) mice at P5. Data are shown as mean±sem, n = 8 mice/group. **p<0.01, ***p<0.001; by nonparametric Mann-Whitney test.

### Increased serum levels of cytokines in mice expressing IL-23 in intestine and skin

To start evaluating the mechanisms leading to stunted growth and early lethality associated with IL-23 expression, we assayed the levels of IL-23 and other cytokines in the serum of *CXR23* and *KSR23* mice. Levels of IL-23 were elevated in the serum of *CXCR3* and *KSR23* mice (~7 and 2.5 fold over controls, respectively) (**Supplemental Fig. 3**). Serum levels of IL-22, IL17, IFN_γ_, IL1β, but not GMCSF, were significantly elevated in the serum of *CXR23* mice (**Supplemental Fig. 3**). Levels of IL-22, IFN_γ_ and IL-1β were significantly elevated in *KSR23* mice (**Fig. 2G**). In general the levels of cytokines in serum correlated with the levels of IL-23, with the changes being more marked in the *CXR23* mice. The most up regulated cytokine in the serum of *CXR23* and *KSR23* mice was IL-22, whose overall levels reached 10 and 4 ng/ml, respectively. The markedly elevated levels of cytokines in serum suggested a possible role for these molecules in pathogenesis.

### Genetic ablation of IL-22 or IL-22R reduces lethality and restores normal body growth in neonates expressing IL-23

To test if IL-22 had a role in the pathogenesis elicited by IL-23 in vivo, we intercrossed *CXR23* and *KS23* mice to IL-22^-/-^ mice generated in our laboratory (**Supplemental Fig. 4**). IL-22-deficient *CXR23 (CXR23/IL-22-*^*/*^*-*) and *KSR23* (KSR23/IL-22^-/-^) mice had extended life spans (**Fig. 3a** and **3e**) and normal growth (**Fig. 3b** and **3f**) suggesting that IL-22 had a major role in the stunted growth and lethality phenotypes exhibited by these animals. Of note, none of the IL-22-deficient *CXR23* mice examined had intestinal bleeding (**Fig. 3c**). To further document a role for IL-22 in pathogenesis we examined animals that still expressed IL-23 and IL-22 but lacked one of the chains of the IL-22R (IL10R2). These mice are referred to as *CXR23/IL10R2*^-/-^ mice. Similar to what was observed above, animals in which IL-22 signaling was inactivated, survived longer (**Fig. 3a**), and had normal body weight at P4 (**Fig. 3b**). Together these results indicate that IL-22 is the main cytokine driving neonatal pathology in mice expressing IL-23 at birth.

**Figure 3 -.**
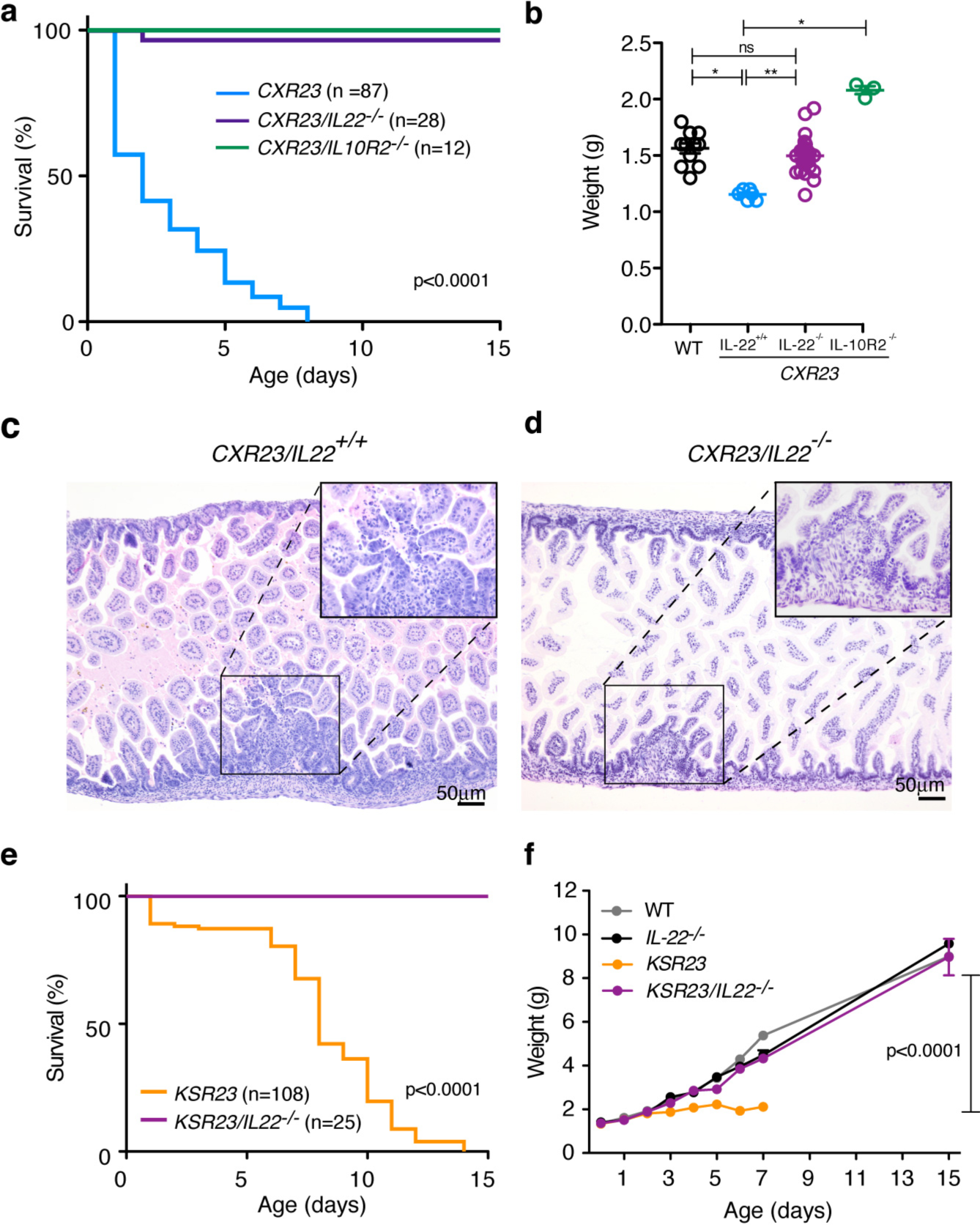
Genetic ablation of IL-22 or IL-22R reduces lethality and restores normal body growth in neonates expressing IL-23. (**a**) Survival curves of *CXR23, CXR23/IL-22*^-/-^ and *CXR23/IL10R2*^-/-^ mice. (**b**) Body weight of WT, *CXR23, CXR23/IL-22’*^*1*^*’*, and *CXR23/IL10R2*^-/-^ at P3-P4 (n= 3-24 mice/group). Representative H&E stained sections of the small intestine of (**c**) *CXR23* and (**d**) *CXr23/IL-22*^-/-^ mice. Inset shows higher magnification of areas with erosive lesions and lymphoid aggregates. (**e**) Survival curves of *KSR23*, and *KSR23/IL-22*mice. (**f**) Weight curves of WT, IL-22^-/-^, *KSR23*, and *KSR23/IL-22*^-/-^ mice over time (n=5-33 mice/group). Data are shown as mean±sem, *p<0.05, **p<0.01; by Mann-Whitney test.

### Increased expression of IL-23 and other cytokines affects expression of genes in neonatal intestine and pancreas

IL-23 and other cytokines were detected at higher levels in the serum of *CXR23* and *KSR23* mice than WT mice. We reasoned that these changes could have affected the growth of the newborn mice. Body growth in newborns is dependent on their ability to ingest, process, and absorb nutrients. Animals in both strains of transgenic mice had milk in the stomach, suggesting that food ingestion was not affected. To investigate if elevated cytokines could affect body growth, we examined the transcriptome of the pancreas (**Fig. 4**) and the intestine (**Fig. 5**), organs that are critical for the processing and absorption of nutrients. To do so we extracted RNA from the pancreas and jejunum of 5 day-old mice expressing IL-23 in skin (*KSR23* mice) and performed RNASeq.

**Figure 4 -.**
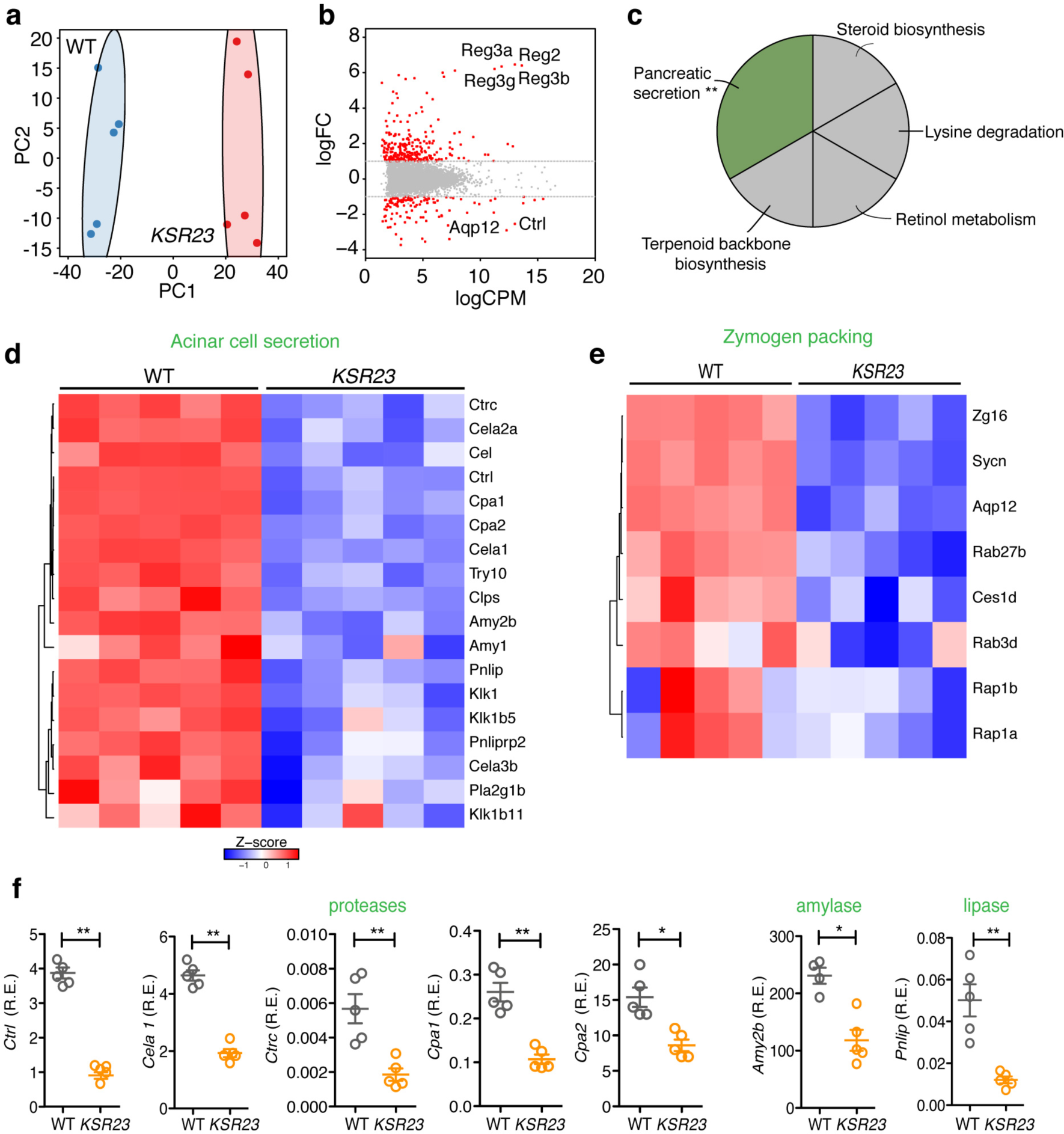
Decreased pancreatic secretion pathway in the pancreas of *KSR23* mice. (**a**) Principal component analysis of RNA-seq expression data from all biological replicates of WT and *KSR23* pancreas (n=5/group). (**b**) Plot of log2FC (log2 fold change) versus log2CPM (log2 counts per million) of all detected transcripts. Points are colored according to expression status: non-significant genes (**gray**) and significant genes (385 genes; Q < 0.05; Log2FC >1 or Log2FC <-1; red). (**c**) Kegg pathway analysis of significantly down regulated genes in the pancreas of *KSR23* mice. (Q<0.05; terms >3 genes; % Genes/term >4; Kappa 0.4). (**d**) Z-scored heat map of downregulated genes associated with the GO terms “pancreatic secretion” and acinar secretory enzymes(Hoang et al., 2016). (**e**) heat mat of downregulated genes associated with the term zymogen packing(Hoang et al., 2016). (**f**) qPCR validation of genes encoding digestive enzymes. Data are shown as mean±sem, *p<0.05, **p<0.01, by nonparametric Mann-Whitney test.

As shown in **Fig 4a** and **4b**, the transcriptome of the pancreas of *KSR23* mice at P5 differed significantly from that of their control littermates. Kegg pathway analysis showed that several genes involved in pancreatic secretion, were downregulated (**Fig. 4c**). The pancreatic secretion pathway includes genes encoding several pancreatic enzymes involved in food digestion and genes involved in the secretory pathway (**Fig. 4c**). Among the genes downregulated in the *KSR23* mice were those encoding pancreatic enzymes, including pancreatic amylases (amylase 2a, *Amy2a;* and amylase 2b, *Amy2b*), lipase (*Pnlip*), elastase (elastase 1, *Cela1; elastase 2a, Cela2a*), and proteases (carboxypeptidase 1, *Cpa 1;* carboxypeptidase 2, *Cpa2;* kallikrein 1-related peptidase b5, *Klk1b5*; chymotrypsin-like protease *Ctrl*; chymotrypsin C, *Ctrc*) (**Fig. 4d**). Other genes involved in pancreatic acinar secretion were also down regulated, including syncollin (*Sycn*), a protein that regulates the fusion of zymogen granules(Wasle et al., 2005). Aquaporin 12 (*AQP12*), an acinar specific water channel that controls secretion of pancreatic fluid(Itoh et al., 2005), was also downmodulated in the pancreas of *KSR23* mice (**Fig. 4e**). Some of these changes were confirmed by qPCR (**Fig. 4f**).

The transcription regulatory gene *Ptf1a* controls acinar cell secretory protein processing and packaging(Hoang et al., 2016). Inactivation of *Ptf1a* in adult acinar cells results in decreased acinar cell identity and this coincides with the appearance of the ductal cell markers Sox9 and keratin19(Hoang et al., 2016; Krah et al., 2015). We observed decreased expression of *Ptf1a* and increased expression of *Sox9* in the pancreas of *KSR23* mice (**Supplemental Fig. 5a** and **b**). These results suggest that acinar cells in *KSR23* mice may acquire ductal cell phenotype. To determine if this was indeed the case, we analyzed the expression of KRT19 in the pancreas of WT and *KSR23* mice by immunostaining (**Supplemental Fig. 5c-f**). Increased number of KRT19+ cells was observed in the pancreas of *KSR23* mice when compared to WT mice (Supplemental Fig 5c and e). KRT19 expression mainly localized to the large interlobular ducts in WT pancreas (**Supplemental Fig 5e**). However, in KSR23 mice, KRT19 also marked a population of acinar cells (**Supplemental Fig. 5f arrow**). These results further indicate that acinar cell function and identity are disturbed in the pancreas of *KSR23* mice.

Trypsinogens, chymotrypsinogens, lipases, elastases and proteases are produced by the acinar cells and secreted via the pancreatic ducts into the small intestine. At this site, some of these precursors are converted into active forms, by enzymes produced by the small intestine, leading to further breakdown of food. Peptides, amino acids, fatty acids, glycerol, and glucose reach the blood stream via transporters present in the brush border membrane of intestinal cells. To ask if increased circulating levels of IL-23 and other cytokines were also associated with marked changes in the expression of genes in the intestine, we analyzed extracted RNA from the jejunum of *KSR23* and control mice and performed RNASeq. The overall pattern of gene expression differed significantly between *KSR23* and control mice (**Fig. 5a**). Several genes were differentially expressed, including IL-22, and genes that are downstream of it such as Reg3 genes (**Fig 5b**) and digestive enzymes also produced by the jejunum. Kegg pathway analysis of WT and *KSR23* jejunum indicated that genes involved in protein digestion and absorption such as chymotrypsinogen B1 (*Ctrbl*), trypsinogen (*Prss1*), trypsin (*Tyr5* and *Tyr4*), caboxypeptidase a1 (*Cpa1*) and chymotrypsin-like elastase family member 3b (*Cela3b*), were downregulated in the intestine of *KSR23* mice (**Fig. 5c**). Transcriptome analysis also showed decreased expression of several genes involved in the regulation of the very low-density lipoprotein particle pathway (VLDL) (**Fig. 5d** and **5e**). These particles regulate fat and cholesterol release into the bloodstream. Together, these results indicate that systemic expression of IL-23, and other cytokines, correlated with significant transcriptional changes in genes that regulate food processing in the pancreas and intestine.

**Figure 5 -.**
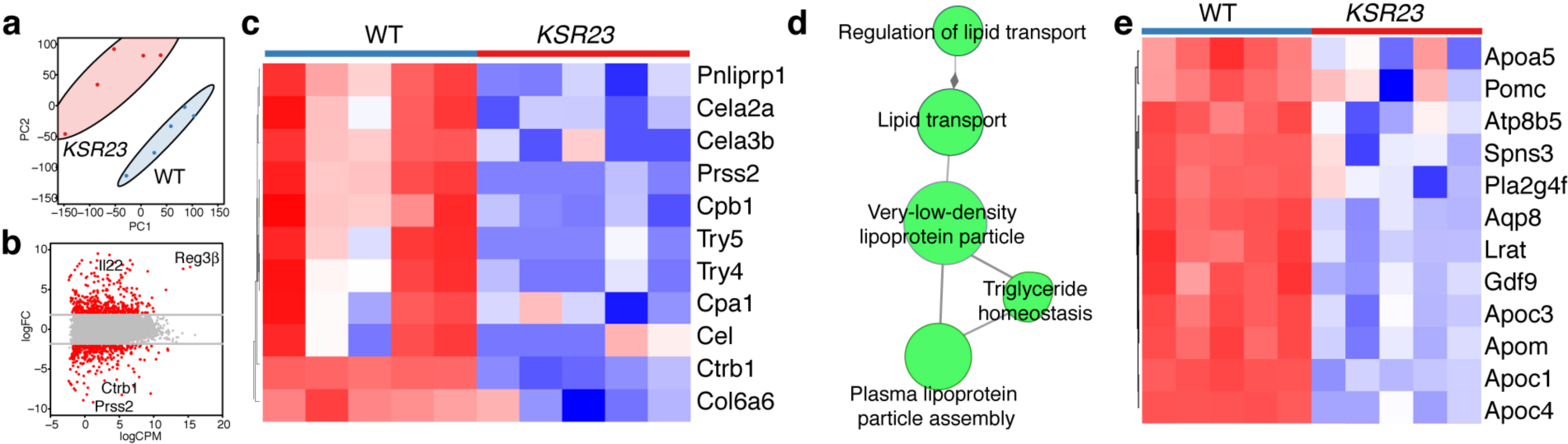
Altered food absorption pathways in the jejunum of *KSR23* mice. (**a**) Principal component analysis of RNA-seq expression data from all biological replicates of WT and *KSR23* jejunum (n=5/group). (**b**) Plot of log2FC (log2 fold change) versus log2CPM (log2 counts per million) of all detected transcripts. Points are colored according to expression status: non-significant genes (**gray**) and significant genes (698 genes; Q < 0.05; Log2FC >2 or Log2FC <-2; red). (**c**) Z-scored heat map of all downregulated genes associated with the GO terms “protein digestion and absorption”. (**d**). ClueGo analysis of significantly down regulated genes shown enrichment of GO term group “very-low-density-lipoprotein-particle” (**e**). Z-scored heat map of all downregulated genes associated with the GO group “very-low-density-lipoprotein-particle”.

### IL-22 ablation corrects aberrant pancreatic enzyme gene expression observed in *KSR23* mice

To ask if IL-22 affected the expression of pancreatic enzymes we examined the expression of several genes by in the pancreas of *KSR23* and *KSR23/IL-22*^-/-^ mice. Quantitative PCR confirmed that *KSR23* mice had increased mRNA levels of Reg3β and decreased levels of pancreatic lipase, amylase and syncollin in the pancreas than that in WT controls (**Fig. 6a, c, e, and g**, respectively). Changes in protein levels were also documented by immunohistochemistry (Fig. 6b, d, f, and h). Deletion of IL-22 in *KSR23* mice had a marked effect on the levels of expression of these genes (**Fig. 6a, c, e, and g**), as well as on the levels of pSTAT3, a known mediator of IL-22 signaling (**Supplemental Fig. 6**). Levels of Reg3β, amylase and pancreatic lipase, as well as syncollin in the pancreas of *KSR23/IL-22*^-/-^ mice were equivalent to those observed in WT mice (**Fig. 6a, c, e, and g**), suggesting that IL-22, or genes downstream of it, had a major role in controlling expression of pancreatic genes.

**Figure 6 -.**
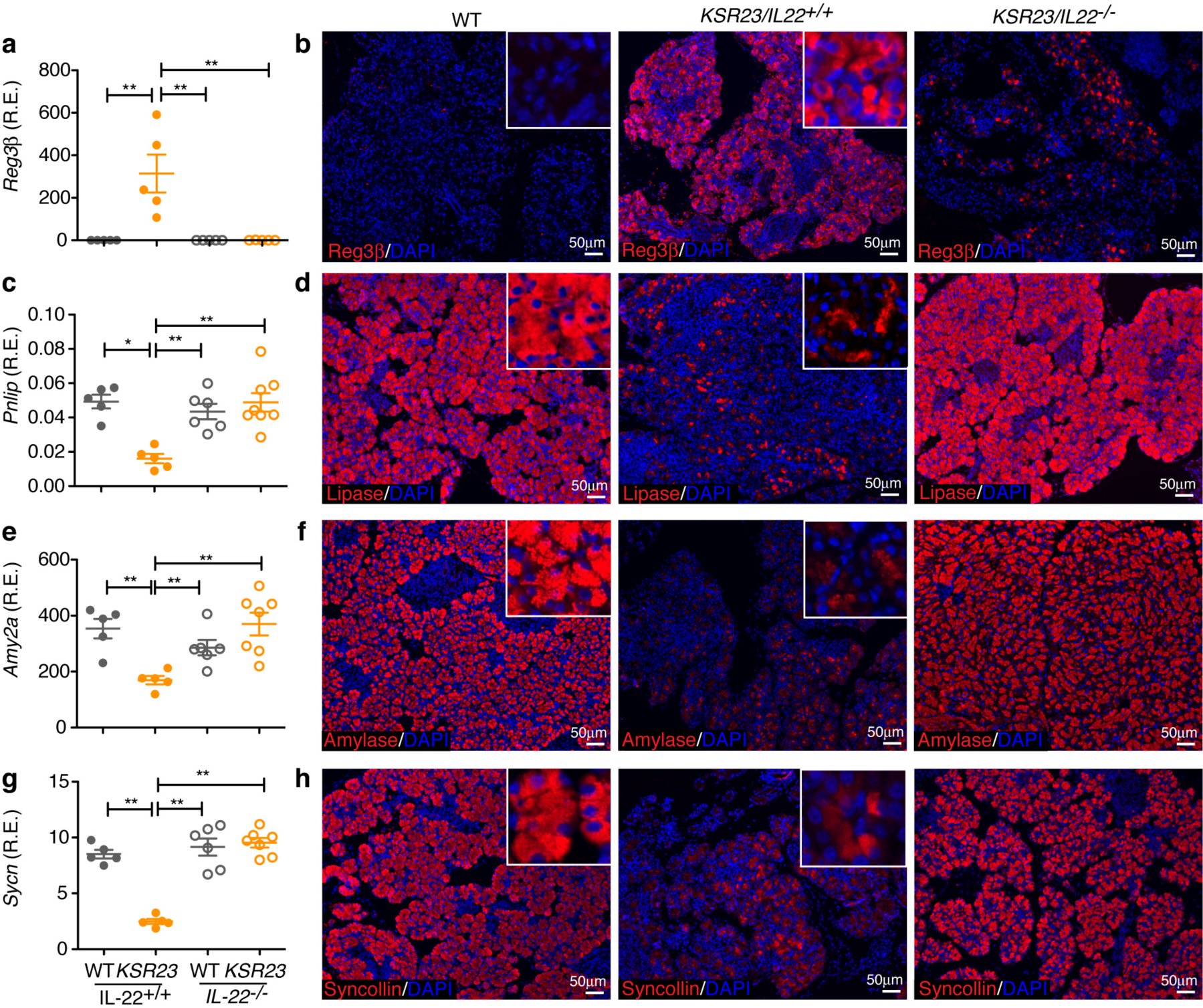
Ablation of IL-22 restores expression of pancreatic genes in *KSR23* mice. (**a, c, e, g**) q-PCR analysis of the expression of selected genes in the pancreas of WT, *IL-22*^-/-^, *KSR23, and KSR23/IL-22*^*/-*^ mice at P5. (b, d, f, h) Immunostaining of pancreas of WT, *KSR23*, and *KSR23/IL-22*^-/-^ mice at P5 with antibodies against Reg3β, lipase (**d**), amylase (**f**), and syncollin (**h**). Insets show higher magnification. Data are shown as mean±sem, n = 5-7 mice/group. *p<0.05, **p<0.01; by nonparametric Mann-Whitney test.

### IL-22 directly regulates expression of pancreatic genes

Deletion of IL-22 reverts the two main phenotypes of early death and stunted body growth in animals expressing IL-23. In addition, deletion of IL-22 reverted the abnormalities in gene expression observed in the pancreas. IL-22R is expressed by acinar cells in the pancreas of WT mice, and it has been shown that IL-22 can directly promote expression of Reg3β by these cells(Aggarwal et al., 2001). To investigate if IL-22 could directly affect expression of pancreatic genes involved in nutrient processing, we performed *in vitro* experiments. Short term (24h) incubation of pancreatic cells with IL-22 led to a significant upregulation of Reg3β as reported(Aggarwal et al., 2001) but no changes in expression of *Ptf1a, Sycn, Amy2a* and *Amy2b* (**Supplemental Fig. 7**). Longer incubation of the acinar cells with IL-22 (60h) led to a significant decrease in the expression of *Ptf1a* (**Fig. 7a**). Coincident with the decreased expression of *Ptf1a* we observed a significant decrease in the expression of *Sycn, Amy2a* and *Prss2* in cultures stimulated with IL-22 (**Fig. 6**), suggesting a direct role for IL-22 in the regulation of these pancreatic genes. To confirm that these changes were elicited by IL-22 signaling, we studied the effect of IL-22 in pancreatic cells derived from IL10R2-deficient mice. No changes in gene expression were observed after addition of IL-22 to IL-22R signaling-deficient pancreatic acinar cells (**Fig. 7** and **Supplemental Fig. 7**), confirming that the changes observed in the expression of pancreatic genes were mediated by the IL-22 receptor. Together these results indicate that the inhibition of pancreatic gene expression observed in *KSR23* mice is mediated by IL-22.

**Figure 7 -.**
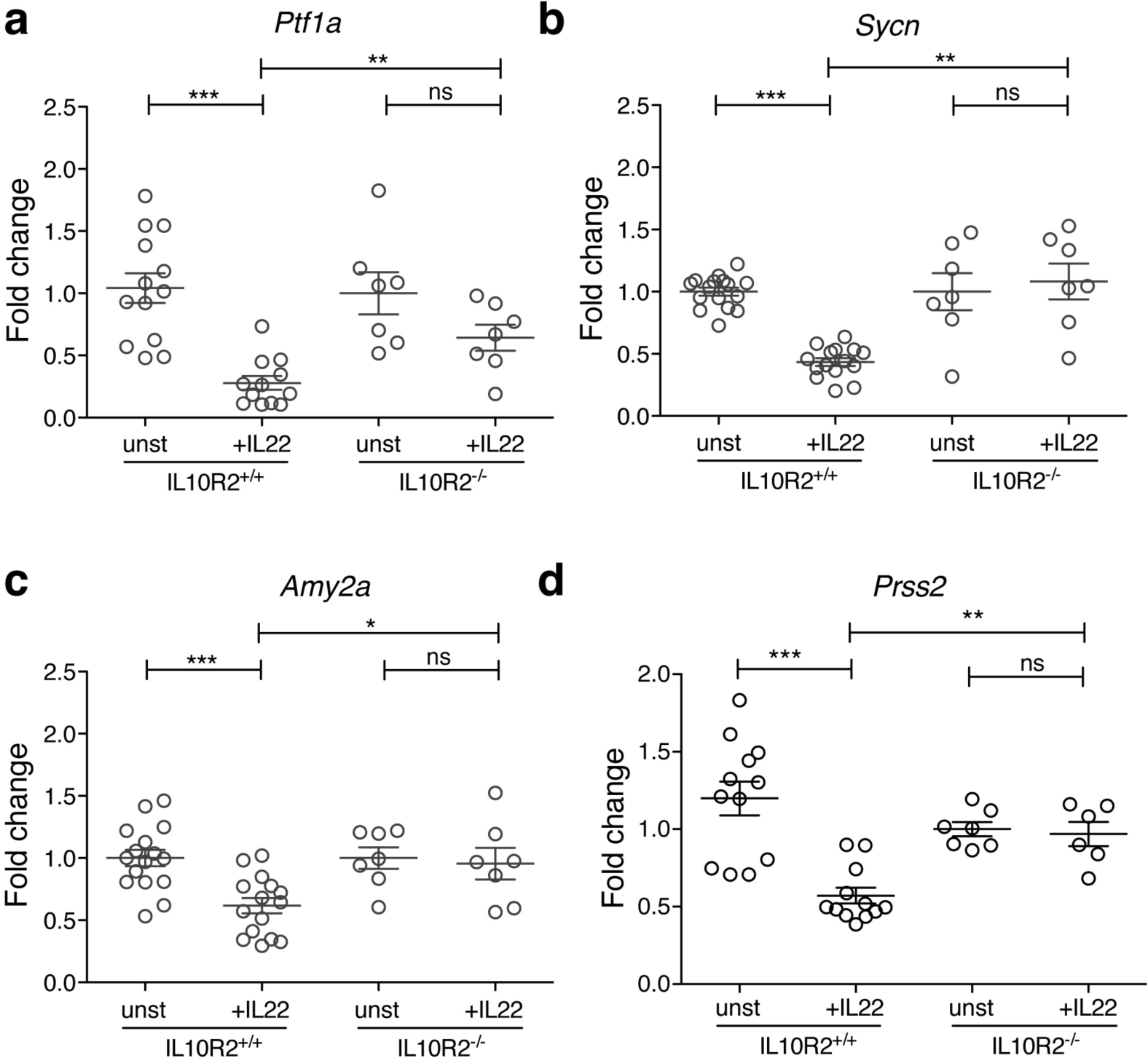
IL-22 induces downregulation of pancreatic enzymes related genes in acinar cell cultures *in vitro* -. Acinar cells from *IL10R2*^*+/+*^ and *IL10R2*mice were cultured in the presence of IL-22 recombinant protein. Sixty hours after culture, cells were analyzed for the expression of (**a**) pancreas transcription factor 1a (*Ptf1a*), (**b**) syncollin (*Sycn*), (**d**) amylase 2a (*Amy2a*), and (**d**) serine protease 2 (*Prss2*) by qPCR. Notice the downmodulation of several pancreatic related genes by IL10R2^+/+^ acinar cells cultured with IL-22. IL-22 did not reduce the expression of pancreatic related gens in IL10R2^-/-^ cells. Data are shown as mean±sem, n= 5-7 mice/group. *p<0.05, **p<0.01, ***p<0.001; by nonparametric Mann-Whitney test.

## Discussion

Here we study the impact of IL-23 expression in newborns. We report that IL-23 induces severe phenotypes when expressed since birth by CX3CR1+ cells and by keratinocytes. These phenotypes are reversed by inactivation of IL-22, suggesting that IL-22 mediates the pathogenic properties of IL-23 in newborns. We also show that a critical and unappreciated pathogenic mechanism triggered by IL-22 is the reduction in the expression of several pancreatic enzymes, which contributes to the failure to thrive observed in these animals.

Two phenotypes were associated with neonatal expression of IL-23. The first was death within the first 48 h of life. This phenotype was present in 50% of the mice expressing IL-23 in CX3CR1+ myeloid cells in the intestine, but not in mice expressing IL-23 in keratinocytes, and was associated with intestinal bleeding, resembling what is observed in animals expressing IL-23 from the villin promoter (V23 mice)(Chen et al., 2015). The phenotype was less severe in the case of *CXR23* mice, perhaps reflecting lower levels of IL-23 expression compared with *V23* mice. The cause of death in both cases is not clear, but the microbiota could play an important role, as germfree *CXR23* mice do not present intestinal bleeding and survive beyond the neonatal period. Also similar to what is observed in the *V23* mice(Chen et al., 2015), we observed an increase in the number of ILC3 in the intestine of *CXR23* mice. We suggest that this cell population is the key driver of pathogenesis in *CXR23* mice, similar to what has been demonstrated in the *V23* mice. Given that ILC3 produce copious amounts of IL-22 upon IL-23 stimulation, it is likely that they are a major source of IL-22 in the newborn gut. Increased IL-22 levels could disrupt intestinal permeability, favor permeation of bacteria into the lamina propria, and cause marked intestinal inflammation and intestinal bleeding and death(Chen et al., 2015). This hypothesis is supported by our observation that *CXR23 germ-free* mice survive beyond the first 48 h, and that IL-22 ablation rescues the early death phenotype.

The second phenotype observed in both strains was failure to growth. *CXR23* mice that survived the first 48 hours failed to grow and died prematurely, similar to *KSR23* mice. Again, these phenotypes were reversed by IL-22 ablation, indicating a pathogenic role for elevated levels of IL-22. Our results are in agreement with other findings in the literature. Transgenic expression of IL-22 from the EμLCK promoter or from the insulin promoter, results in animals with small body weight that die a few days after birth(Wolk et al., 2009). Expression of IL-22 from the albumin promoter, on the other hand, does not induce early lethality, but the body weight of the animals is lower than that of wild type littermates from 5 months of age on(Park et al., 2011). The reason why animals in this line survive, and the other strains driven by the EμLCK and insulin(Wolk et al., 2009), villin(Chen et al., 2015) and CX3CR1 promoters do not, is unclear, but it might be related to overall levels of IL-22 in circulation or the site and timing of its expression. Low body weight is also observed in mice in which IL-22 was delivered by adenoviral infection or in animals injected daily with this cytokine (25 mg over a 2-wk period). Daily injection of IL-22 induced a 8% decrease in body weight(Liang et al., 2010). The causes of stunted growth and low body weight are multiple and include reduced absorption of dietary components. Indeed, it was suggested previously that IL-22 regulates expression of lipid transporters in the intestine(Mao et al., 2018), a finding that we confirm and extend here.

The most unexpected finding in the course of our experiments was that increased levels of IL-22 in circulation markedly affected the expression of pancreatic enzymes critical for digestion of carbohydrates, proteins and lipids. The reduced expression of enzymes and reduced absorption of nutrients by the intestine resulted in malabsorption, as evidenced by increased amount of fat in the stools. The expression of the digestive enzymes was normalized by deletion of IL-22. Thus, it is likely that the growth disturbances observed in the *CXR23* and *KSR23* mice result in part from malabsorption of critical dietary components due to reduced expression of pancreatic enzymes and of intestinal transporters.

While we cannot rule out that the other cytokines simultaneously upregulated by IL-22 may impact the expression of the pancreatic and intestinal genes, our demonstration that IL-22 can directly downregulate expression of genes encoding the pancreatic enzymes *in vitro*, suggest that IL-22 is the main factor regulating this process in IL-23 expressing animals. We show here that incubation of acinar cells with IL-22 leads to a significant reduction in the expression of *Ptf1a.* Ptf1a is a basic helix-loop-helix transcription factor that is critical for the specialized phenotype of the pancreatic acinar cells, including the control of the production of secretory digestive enzymes(Krah et al., 2015). We show an increase in the number of pSTAT3 acinar cells of the *KSR23* mice and postulate that this may be the effector mechanism for the reduced expression of digestive enzymes. Interestingly, the genetic ablation of Ptf1a in adults leads to reduction in the expression of genes encoding acinar secretory products and genes involved in the zymogen granule formation(Hoang et al., 2016). Also of interest is the observation that decreased Ptf1a expression results in increased expression of ductal cell markers(Hoang et al., 2016; Krah et al., 2015). Similar results were observed in *KSR23* mice. Our observations linking high systemic levels of IL-22 with reduced levels of Ptf1a expression in the pancreas of *KS23* mice, and the demonstration of a direct activity of IL-22 on the acinar cell, strongly support a role for IL-22 in the expression of *Ptf1a.*

The IL-22-induced upregulation of several antimicrobial genes by acinar cells is yet another interesting observation in this context. IL-22 directly regulates expression of Reg3β *in vitro*, as shown here and elsewhere(Aggarwal et al., 2001). The additional secretion of the microbicide Reg proteins into the intestinal lumen could potentially affect the intestinal microbiota. Thus, the fact that IL-22 can regulate expression of several pancreatic genes, may have implications for the pathogenesis of other diseases of the intestinal tract. Varying degrees of pancreatic insufficiency have been reported in patients with inflammatory bowel disease (IBD) (Barthet et al., 1999; Barthet et al., 2006; Bokemeyer, 2002), and these may be functionally related to higher levels of IL-22 in situ or in circulation. Relevant to this discussion is the observation that pediatric IBD is often associated with growth failure(Diefenbach and Breuer, 2006), which is caused by several factors among them decreased food intake and malabsorption(Rosen et al., 2015).

In summary, we have demonstrated an important role for IL-23 in neonatal pathology. The main effector mechanism triggered by IL-23 is increased production of IL-22, which acts at the level of the intestinal epithelium and acinar cells in the pancreas to regulate intestinal permeability and inhibit mechanisms associated the digestion and absorption of nutrients. These results indicate that dysregulation in the expression of IL-23 and IL-22 has a significant negative impact on survival and overall fitness of newborns.

## Methods

### Mice

*R23* mice were described in(Chen et al., 2018). *R23* mice were backcrossed into the C57BL/6 background for 11 generations. Generation of IL-22-deficient mice in the C57BL/6 background is described in **Supplemental Fig. 4**. C57BL/6 (CN 000664), CX3CR1-cre(Yona et al., 2013) (CN 025524), hK14-cre(Dassule et al., 2000) (CN 018964), and IL10R2K0(Spencer et al., 1998) (CN 005027) mice were purchased from The Jackson laboratory (Bar Harbor, ME). IL-22 tDt-tomato mice were provided by Dr. Scott Durham (NIH, Bethesda, MD). Mice were maintained under specific pathogen-free conditions. All experiments involving animals were performed following guidelines of the Animal Care and Use Committee of the Icahn School of Medicine at Mount Sinai.

### Generation of IL-22-deficient mice

IL-22 deficient (*IL-22*^-/-^) mice were generated using CRISPR/Cas9 technology directly into C57BL/6 as described before(Mashiko et al., 2013). To do so, we designed a sgRNA that targeted IL-22 exon 1 by using the online CRISPR Design Tool (http://tools.genome-engineering.org). The plasmid expressing sgRNA was prepared by ligating oligos into BbsI site of pX330 (Addgene plasmid ID: 42230) (Ran et al., 2013). Confirmation of WT and KO genotypes was done by sequencing. Functional validation of the IL-22 knockout was done as described in **Supplemental Fig. 4**.

### Flow cytometry

The small intestine (SI) of P1 mice were microdissected using a stereo microscope, minced and digested with 2 mg ml^-1^ collagenase D (Roche). The cell suspension was passed through a 70µm cell strainer and mononuclear cells were isolated. Cells were preincubated with anti-mouse CD16/CD32 for blockade of Fc receptors, washed and incubated for 40 min with the appropriate monoclonal antibody conjugates in a total volume of 200 μ! PBS containing 2 mM EDTA and 2% (vol/vol) bovine serum. Propidium iodide (Sigma Aldrich) or DAPI (Invitrogen) was used to distinguish live cells from dead cells during cell analysis. Stained cells were analyzed on a FACS Canto or LSRII machine using the Diva software (BD Bioscience). Data were analyzed with FlowJo software (TreeStar). ILC3 cells were defined as CD45^+^Lin^-^Thy1^+^Sca-1^hi^ as described(Chen et al., 2015).

### Enzyme-linked immunoabsorbent assay

Chemokines and cytokines (GCSF, GMCSF, CXCL1, CXCL2, CXCL10, IFN_γ_, IL1a, Μβ, IL12p70, IL15/IL15R, IL17, IL18, IL21, IL-22, IL-23, IL25, IL27, Leptin, CCL2, CCL3, CCL4, CCL5, CCL7, MCSF, sRankL, TNF) were measured in mouse serum with ProcartaPlex Multiplex Immunoassays (eBioscience) according to the manufacturer’s protocol. Analysis was performed with the xMAP Technology by Luminex.

### Triglycerides

Blood was collected from WT and *KSR23* mice at P5 after a 4 h starvation period. Serum separation was performed using the Capillary Blood Collection Tubes (Terumo™ Capiject™). Serum triglyceride levels were measured with the Triglyceride (GPO) reagent set (Pointe Scientific, INC) according to the manufacturer’s specifications. To measure triglycerides in the stool we collected 1 mg of feces in a 1.5 ml eppendorf tube and resuspended it in 500 μl of phosphate buffered saline (PBS). 500 μl of chloroform in methanol solution (2:1) was added to the fecal suspension and vortexed. The suspension was centrifuged (1000 x g, 10 min, at room temperatre) and the lower liquid phase, containing the extracted lipids in chloroform:methanol, was transferred to a new 1.5 mL Eppendorf tube. Samples were incubated for 24 h at 37°C in order to evaporate all liquid. The lipid pellet was then resuspended in 200 uL PBS and the triglyceride concentration measured with the Triglyceride (GPO) reagent set (Pointe Scientific, INC) according to the manufacturer’s specification.

### Acinar cell culture

Pancreatic acinar cells were isolated as described(Gout et al., 2013) and cultured for 24 and 60 hours in Waymouth’s medium alone or with addition of recombinant IL-22 (50 ng/mL, Kingfisher Biotech) at 37 °C, 5% CO_2_.

### Reverse-transcription polymerase chain reaction

Total RNA from tissues was extracted using the RNeasy mini/micro Kit (Qiagen) according to the manufacturer’s instructions. Complementary DNA (cDNA) was generated with Superscript III (Invitrogen). Quantitative PCR was performed using SYBR Green Dye (Roche) on the 7500 Real Time System (Applied Biosystems) machine. Results were normalized to the housekeeping gene Ubiquitin. Relative expression levels were calculated as 2^(Ct(Ubiquitin)-Ct(gene))^. Primers were designed using Primer3Plus and are listed in **Supplemental Figure 8**.

### RNA-seq

Small intestine and pancreas were homologized in Trizol reagent (Invitrogen). Total RNA from tissues was extracted using the RNeasy mini Kit (Qiagen) according to the manufacturer’s instructions. Samples were shipped on dry ice to the Center for Functional Genomics and the Microarray & HT Sequencing Core Facility at the University at Albany (Rensselaer). RNA quality was assessed using the Nanodrop (Thermo Scientific) and Bioanalyzer Total RNA Pico assay (Agilent). Total RNA with a RNA integrity number (RIN) value of 8 or greater was deemed of good quality to perform the subsequent protocols. 100 pg of total RNA was oligo-dT primed using the SMART-Seq v4 Ultra Low Input RNA Kit (Clontech) and resulting the cDNA was amplified using 15 cycles of PCR. The double stranded cDNA (dscDNA) was purified using AMPure XP magnetic beads and assessed for quality using the Qubit dsDNA HS assay and an Agilent Bioanalyzer high sensitivity dscDNA chip (expected size ~600bp-9000bp). The Illumina Nextera XT kit was used for library preparation wherein 125 pg dscDNA was fragmented and adaptor sequences added to the ends of fragments following which 12 cycles of PCR amplification was performed. The DNA library was purified using AMPure XP magnetic beads and final library assessed using Qubit dsDNA HS assay for concentration and an Agilent Bioanalyzer high sensitivity DNA assay for size (expected range ~600-740bp). Library quantitation was also done using a NEBNext Library Quant kit for Illumina. Each library was then diluted to 4nM, pooled and denatured as per standard Illumina protocols to generate a denatured 20 pM pool. A single end 75bp sequencing was performed on the Illumina Nextseq 500 by loading 1.8 pM library with 5% PhiX on to a 75 cycle high output flow cell. The RNAseq data was checked for quality using the Illumina FastQC algorithm on Basespace.

### Transcriptome analyses

RNA-Seq data from small intestine and pancreas was mapped to the mouse reference genome (UCSC/mm10) using Tophat version 2.1.0(Trapnell et al., 2009). Gene-level sequence counts were extracted for all annotated protein-coding genes using htseq-count version 0.6.1(Anders et al., 2015) by taking the strict intersection between reads and the transcript models associated with each gene. Raw count data were filtered to remove low expressed genes with less than five counts in any sample. Differentially expressed genes between groups were analyzed using Bioconductor EdgeR package version 3.10.2 Bioconductor/R(McCarthy et al., 2012; Robinson et al., 2010). Statistically significant differentially expressed genes between groups (Q < 0.05) were selected in gene-wise log-likelihood ratio tests that were corrected for multiple testing by Benjamini and Hochberg FDR. KEGG pathway enrichment analyses were performed using ClueGo(Bindea et al., 2009; Shannon et al., 2003) to identify pathways in which differentially expressed genes are involved. A cut-off of 0.4 was set for kappa score and terms including at least 3 genes were retrieved.

### Immunostaining and imaging

Organs were dissected, fixed in 10% phosphate-buffered formalin, and processed for paraffin sections. Fourµm sections were dewaxed by immersion in xylene (twice for 5 minutes each time) and hydrated by serial immersion in 100%, 90%, 80%, and 70% ethanol and PBS. Antigen retrieval was performed by microwaving tissue sections for 15 minutes in Target Retrieval Solution (DAKO). Tissue sections were incubated with primary Abs in a humidified atmosphere for 1 hour at room temperature. After washing, conjugated secondary Abs were added and then incubated for 35 minutes. The slides were then washed and mounted with Fluoromount-G (SouthernBiotech). Primary and secondary antibodies used are listed in **Supplemental Fig. 9**. Immunofluorescent imaging was performed in a fluorescence microscope (Nikon Eclipse *Ni*) with Plan Apo objective lenses. Images were acquired using a digital camera (DS-QiMc; Nikon) and Nis-Elements BR imaging software. Images were composed in Adobe Photoshop CS3.

### Statistics

Differences between groups were analyzed with Student’s t test or nonparametric Mann-Whitney test. For the comparison of more than two groups a one-way ANOVA followed by a Bonferroni multiple comparison test was performed. Survival curves were analyzed by a log-rank test. All statistical analyses were performed with GraphPad Prism 5 software.

## Author contributions

G.C.F, L.C, V.S, Z.H, M.D, and J.F did experiments and analyzed data. T.H.M and T.K assisted with the Luminex experiments. S.D and H.X provided reagents. G.C.F and S.A.L designed study, analyzed data and wrote the manuscript. All authors reviewed and edited the manuscript.

## Acknowledgments

We thank all members of the Lira Lab for discussions and support. We thank Dr. Maria C Lafaille for comments. We thank Ana Valbuena for help with the Luminex assay. We thank Dr. Kevin Kelley and the Mouse Genetics and Gene Targeting CoRE Facility for assistance in generation of gene-modified mice. L.C. was supported by a Research Fellowship Award from the Crohn’s & Colitis Foundation of America (CCFA). This work was supported by the grant R01 1R01DK110352

---

**Supplemental Figure 1.**
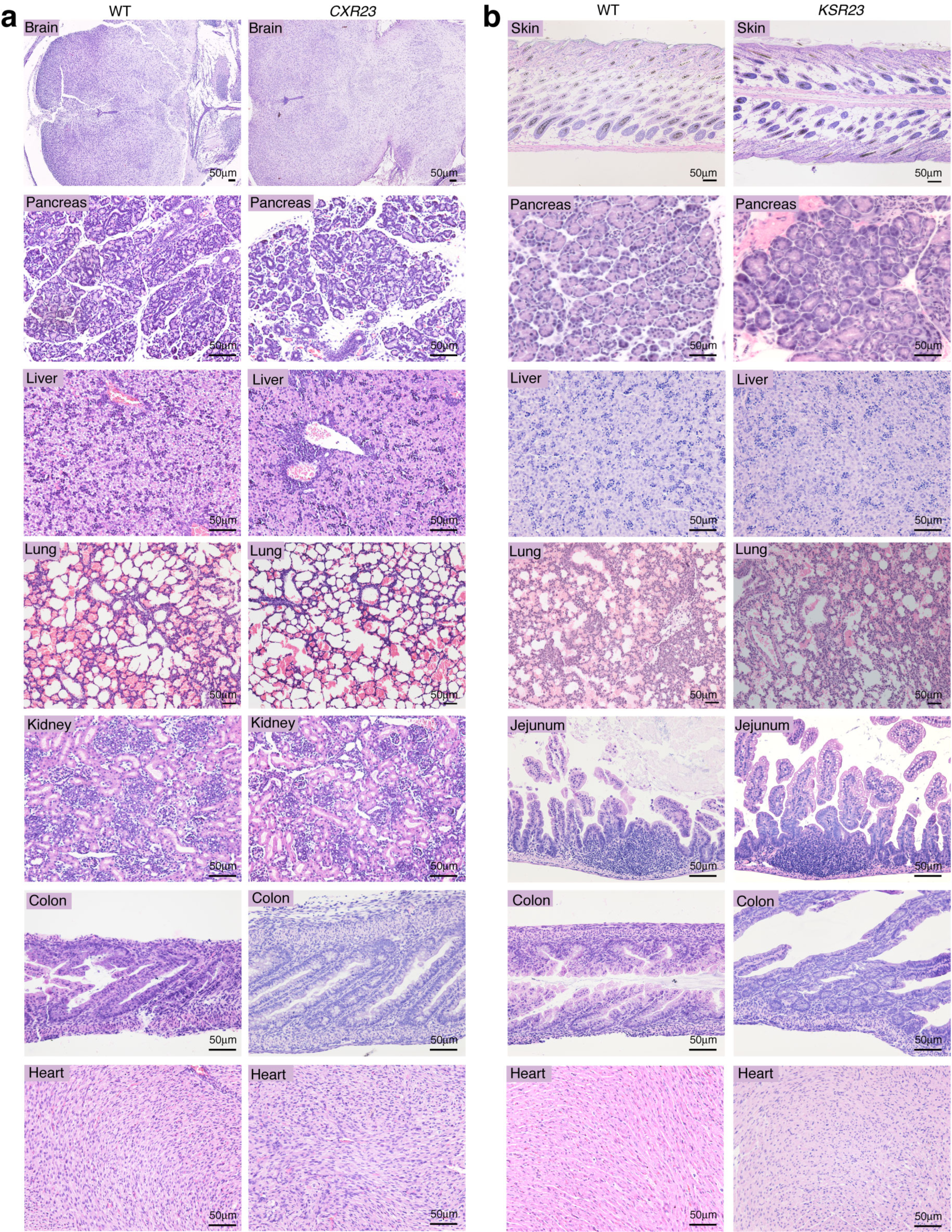
*CXR23* and *KSR23* mice do not develop systemic inflammation. Representative H&E section of indicated organs from (**a**) WT and *CxR23* mice at P1 and (**B**) WT and *KSR23* at P5.

**Supplemental Figure 2.**
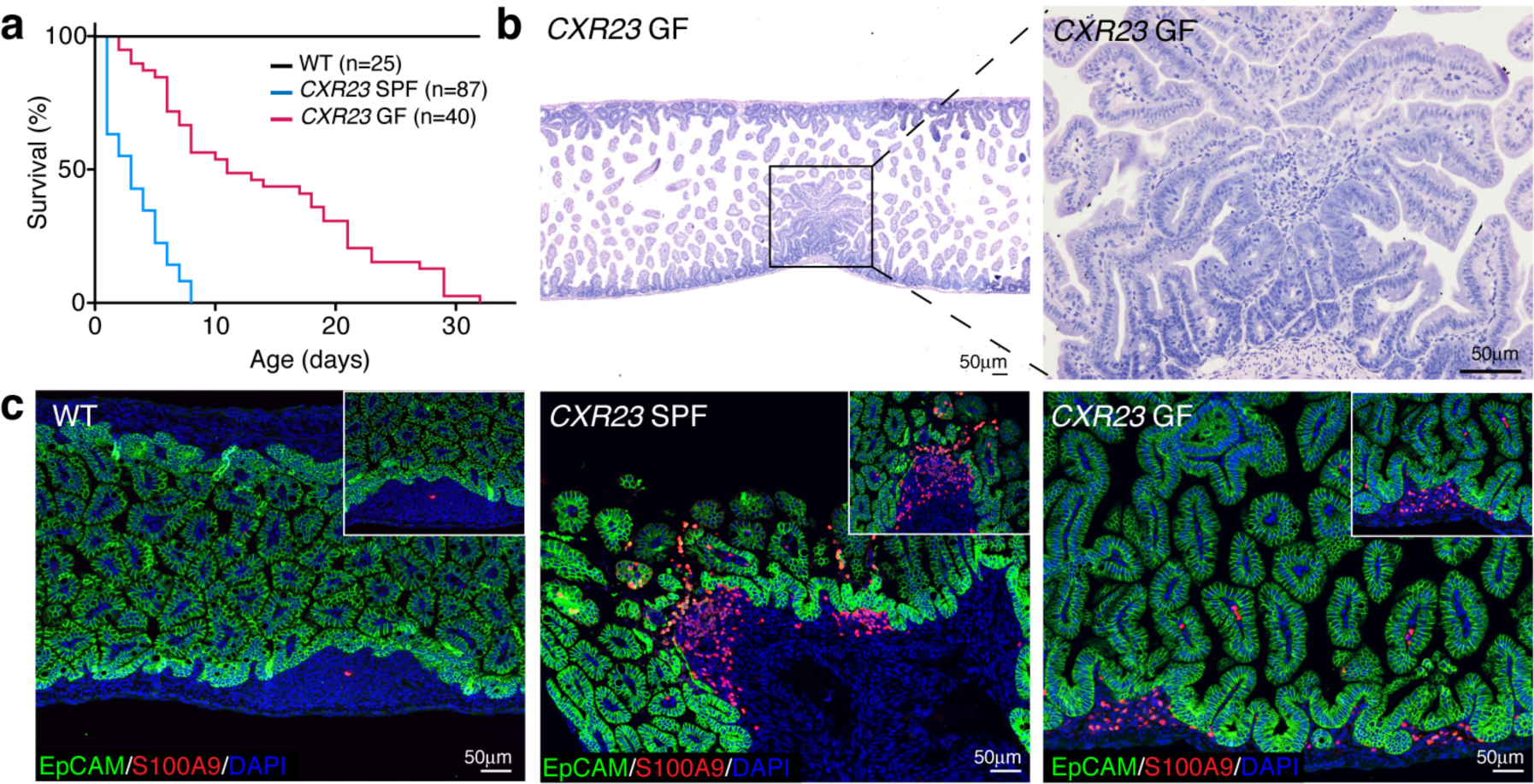
*CXR23* germ-free mice have increased lifespan when compared to *CXR23* SPF mice but die at an early age. (**a**) Survival curves of WT, *CXR23* SPF and *CXR23* GF mice over time. (**b**) Representative H&E section of the small intestine of *CXR23* GF mice at P7. (**c**) Immunostaining of the small intestine of *CXR23* SPF and *CXR23* GF mice showing the accumulation of S100A9^+^ neutrophils. Zoomed-in boxed area shows disrupted EpCAM^+^ intestinal epithelial cells and extravasation of neutrophils into the gut lumen of *CXR23* SPF mice. Neutrophils accumulate in areas contining intact epithelium in *CXR23* GF mice (inset).

**Supplemental Figure 3.**
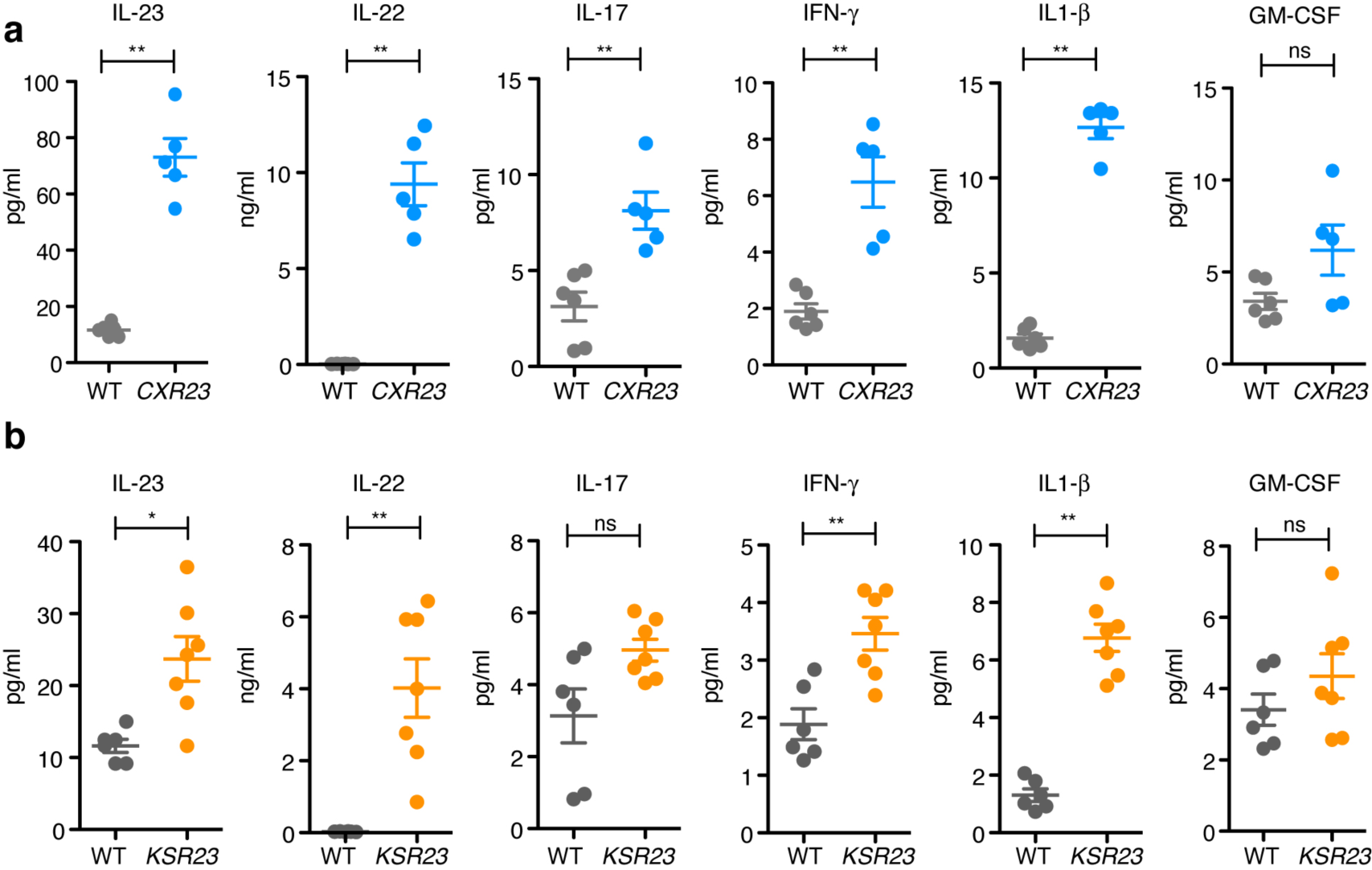
Quantification of cytokines in the serum of WT (P1-P5), (**a**) *CXR23* mice at P1 and (**b**) *KSR23 mice at* P5 (n =5-8 mice/group). Data are shown as mean±sem, *p<0.05, **p<0.01, ***p<0.001; by nonparametric Mann-Whitney test.

**Supplemental Figure 4.**
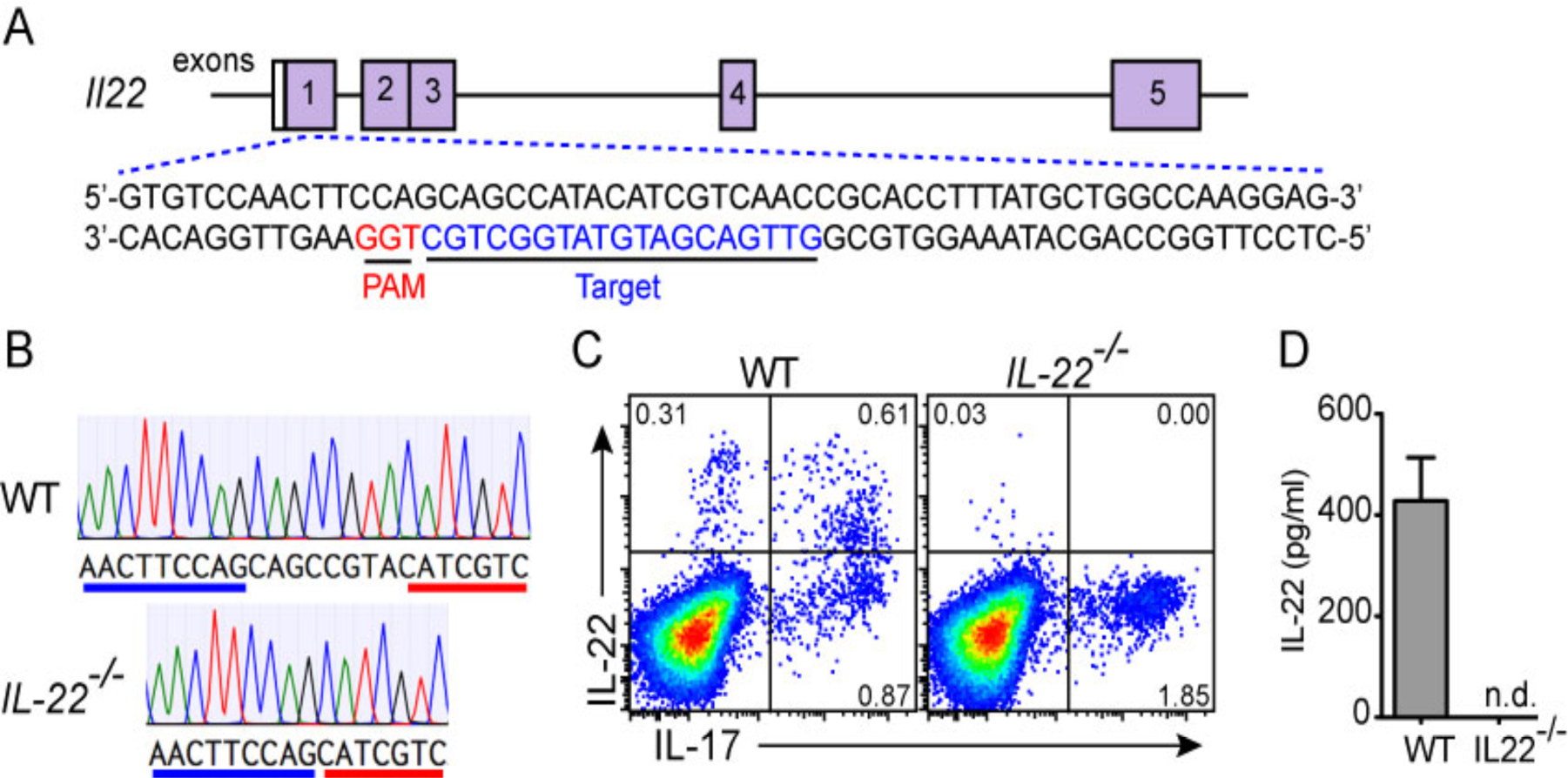
Generation of IL-22-deficient mice by CRISPR/Cas9 technology. (**a**) Schematic illustration of the IL-22 gene structure and part of exon 1 sequences. The target sequence and PAM domain are indicated in blue and red, respectively. (**b**) Genomic sequences of IL-22 (exon 1) from the WT and *IL-22*^-/-^ mice. Note a 8bp nucleotide deletion in the *IL-22*^-/-^ mice, which causes frame-shifting and generates a premature stop codon. (**c**) FACS intracellular staining of IL-22 and IL-17 in IL-23 activated CD4^+^ T cells. (**d**) IL-22 levels measured by ELISA in the supernatant of activated CD4^+^ T cell cultures. n.d., undetectable. n=3 mice per group.

**Supplemental Figure 5.**
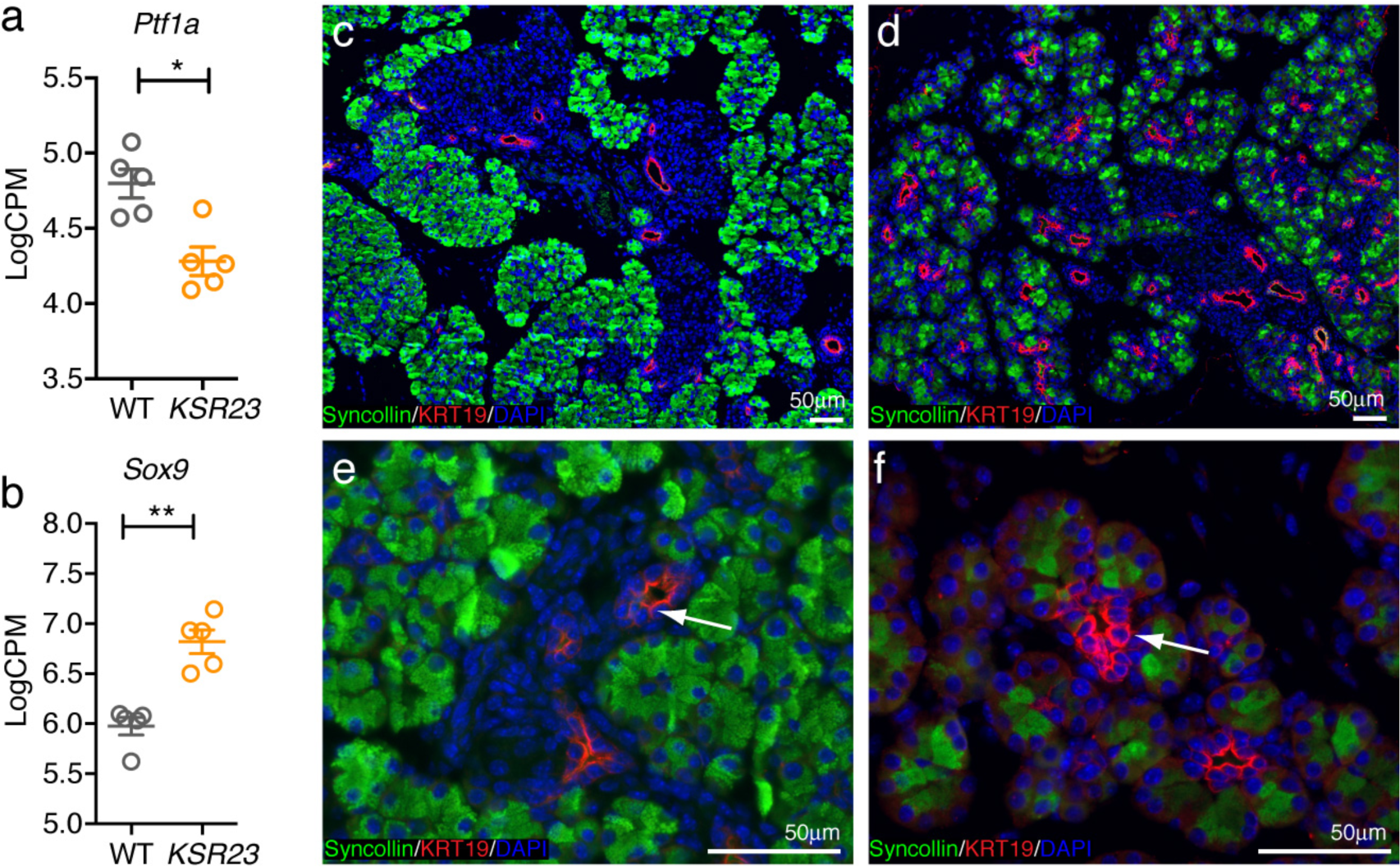
Increased number of ductal cells in the pancreas of *KSR23* mice. RNAseq analysis of the expression of (**a**) *Ptf1a* and (**b**) *Sox9* in the pancreas of WT and *KSR23* mice at P5 (n=5 mice/group). Immunostaining of pancreas of WT (**c & e**) and *KSR23* (d & e) mice with antibodies against syncollin and KRT19. Arrows show localization of KRT19 positive cells in the pancreas of WT (**e**) and KSR23 (**f**) mice. Data shown as mean±sem,. *p<0.05, **p<0.01; by nonparametric Mann-Whitney test.

**Supplemental Figure 6.**
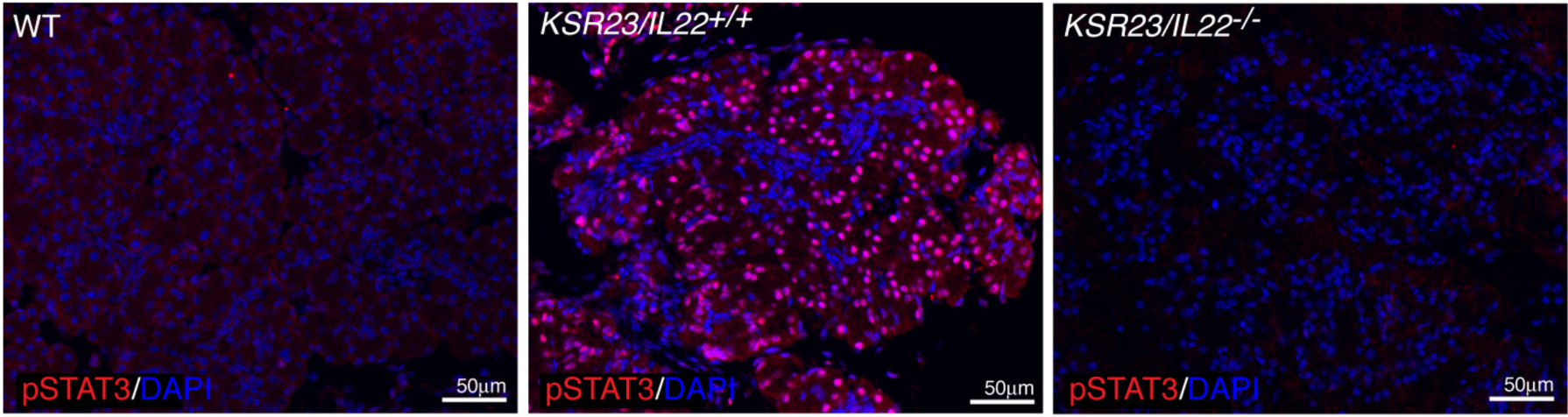
Increased number of pSTAT3^+^ cells in the pancreas of *KSR23* mice. Immunostaining of pSTAT3 in the pancreas of WT, *KSR23 and KSR23/IL-22*^-/-^ mice.

**Supplemental Figure 7.**
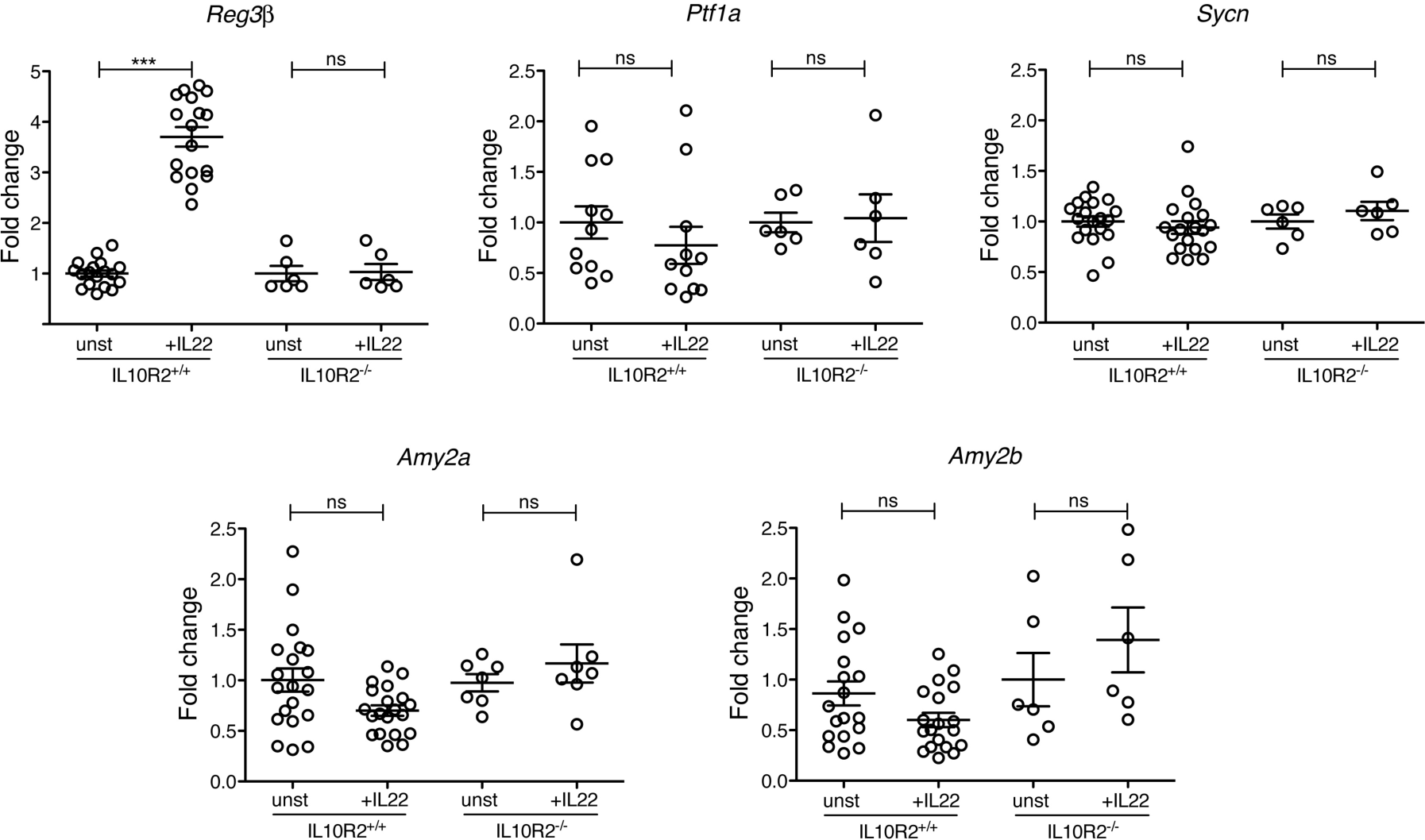
IL-22 induces upregulation of *Reg3β* but does not alter expression of pancreatic enzymes in acinar cell cultures 24h after IL-22 stimulation. Acinar cells from IL10R2^+/+^ and IL10R2^-/-^ mice were cultured in the presence of IL-22 recombinant protein. Twenty-four hours after culture, cells were analyzed for the expression of pancreas transcription factor 1a (*Ptf1a*), syncollin (*Sycn*), amylase 2a (*Amy2a*), and amylase 2b (*Amy2b*) by qPCR. The expression of *Reg3β* was significantly increased at 24 h. However, there were no changes in the expression of other pancreatic related genes by IL10R2^+/+^ cells cultured with IL-22. IL-22 did not reduce the expression of pancreatic related gens by IL10R2^-/-^ cells. Data are shown as mean±sem, n = 5-7 mice/group. ***p<0.001; by nonparametric Mann-Whitney test.

**Supplemental Figure 8.**
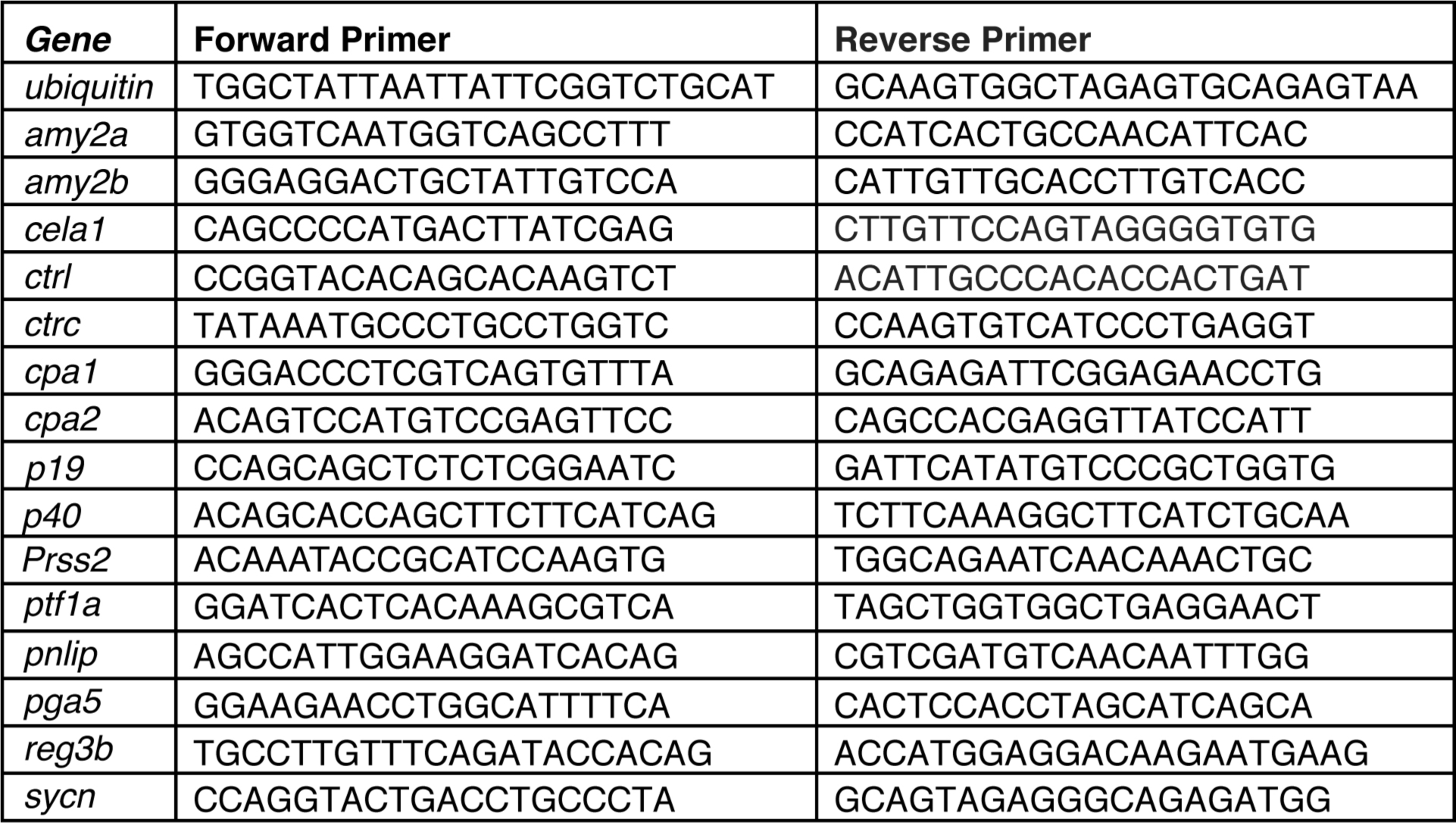
List of primers used in qPCR studies -

**Supplemental Figure 9.**
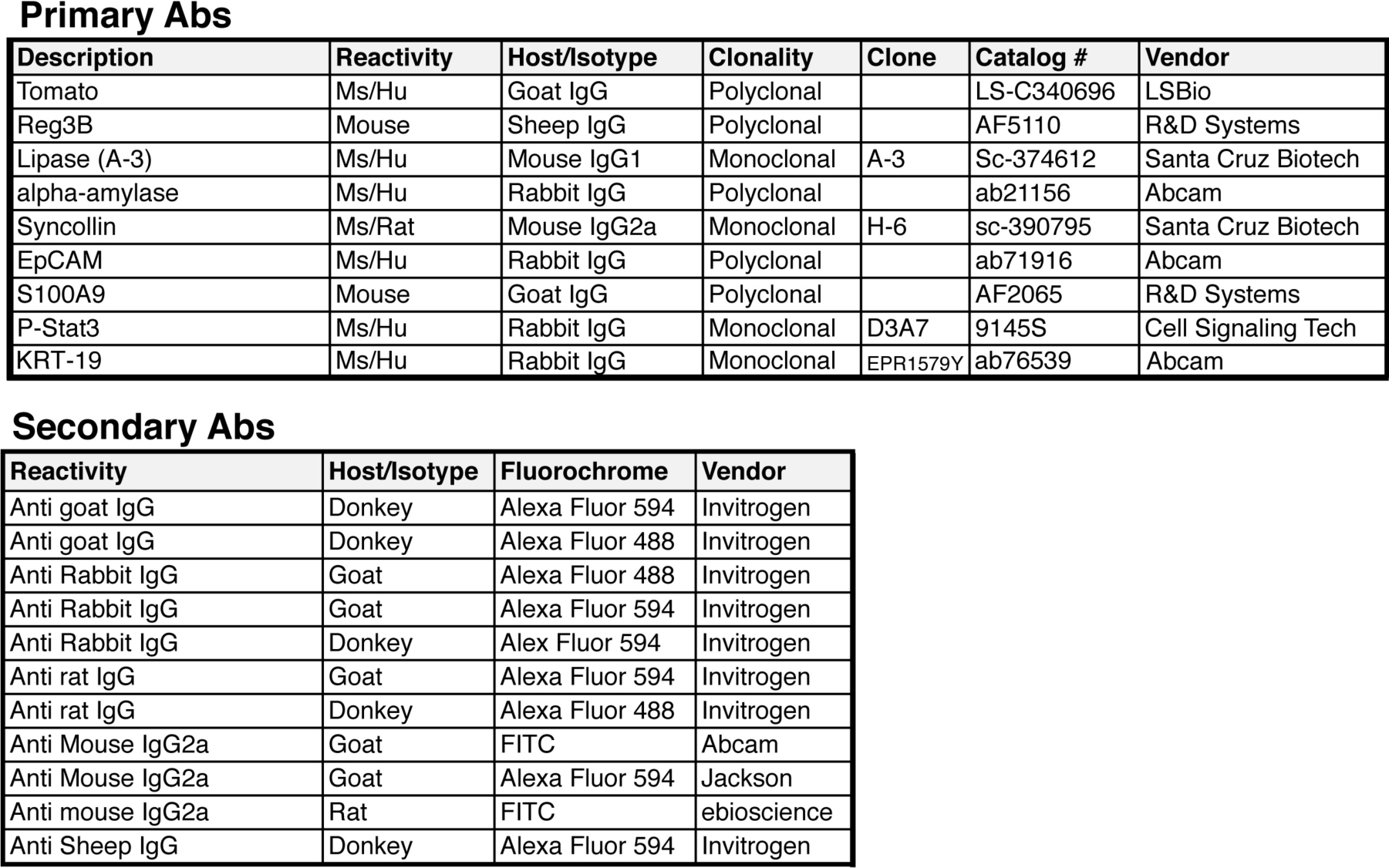
List of antibodies used in immunofluorescence studies -

